# Plant organic matter inputs exert a strong control on soil organic matter decomposition in a thawing permafrost peatland

**DOI:** 10.1101/2021.10.20.465126

**Authors:** Rachel M. Wilson, Moira A. Hough, Brittany A. Verbeke, Suzanne B. Hodgkins, IsoGenie Coordinators, Jeff P. Chanton, Scott D. Saleska, Virginia I. Rich, Malak M. Tfaily

## Abstract

Peatlands are a climate critical carbon (C) reservoir that will likely become a C source under continued warming. A strong relationship between plant tissue chemistry and the soil organic matter (SOM) that fuels C gas emissions is inferred, but rarely examined at the molecular level. Here we compared Fourier transform infrared (FT-IR) spectroscopy measurements of solid phase functionalities in plants and SOM to ultra-high-resolution mass spectrometric analyses of plant and SOM water extracts across a palsa-bog-fen thaw and moisture gradient in an Arctic peatland. From these analyses we calculated the C oxidation state (NOSC), a measure which can be used to assess organic matter quality. Palsa plant extracts had the highest NOSC, indicating high quality, while extracts of *Sphagnum*, which dominated the bog, had the lowest NOSC. The percentage of plant compounds that are less bioavailable and accumulate in the peat, increases from palsa (25%) to fen (41%) to bog (47 %), reflecting the pattern of percent *Sphagnum* cover. The pattern of NOSC in the plant extracts was consistent with the high number of consumed compounds in the palsa and low number of consumed compounds in the bog. However, in the FT-IR analysis of the solid phase bog peat, carbohydrate content was high implying higher quality SOM. We explain this discrepancy as the result of low solubilization of bog SOM facilitated by the low pH in the bog which makes the solid phase carbohydrates less available to microbial decomposition. Plant-associated lignins and tannins declined in the unsaturated palsa peat indicating decomposition, but accumulated in the bog and fen peat where decomposition was presumably inhibited by the anaerobic conditions. A molecular-level comparison of the aboveground C sources and peat SOM demonstrates that climate-associated vegetation shifts in peatlands are important controls on the mechanisms underlying changing C gas emissions.

## Introduction

Climate-change induced warming, especially in the Arctic, will provoke a series of responses including changes to the overall plant community composition, individual plant primary productivity, microbial community, and individual microorganisms which culminate in the whole ecosystem response (Wardle et al., 2004). These interactions are complex and their interpretation is, in turn, complicated by the extreme complexity of the soil organic matter (SOM) that acts as the repository of plant derived substrates, inhibitory compounds, and microbially derived metabolic products. Understanding such interactions is critical because interactions between plants and the microbial community have a strong effect on the net release of the microbial respiration products CO_2_ and CH_4_ (Sutton-Grier and Megonigal 2011).

Peatlands are a globally significant carbon (C) reservoir estimated at 530 ± 160 Pg (Hugelius et al., 2020) up to 1055 Pg (Nichols and Peteet 2019), representing 35-70% of planetary soil organic carbon (Lal 2010). Much of the high-latitude peatland C (277-800 Pg) is currently protected from decomposition as peatland permafrost (Tarnocai et al., 2009; Hugelius et al., 2014). Due to climate change, northern high latitudes are warming two to three times faster than the global average (Rintoul et al., 2018), which is causing permafrost to thaw (Christensen, 2014). Once thawed, the soil organic C is susceptible to microbial decomposition into the potent greenhouse gases (GHG) carbon dioxide (CO_2_) and, under water-logged anaerobic conditions, methane (CH_4_) (Schaedel et al., 2016; Chang et al., 2021). Many peatlands are active C sinks (Turetsky et al., 2007; Jones et al., 2013) or near-C neutral (Zoltai 1993, Euskirchen et al., 2012). The source or sink potential of a peatland depends on the balance between net C uptake by primary production and C loss via heterotrophic respiration, both of which can be affected by climate change. C uptake increases under a longer growing season (Natali et al., 2012), warming, drying (e.g., Malhotra et al., 2020) and changing plant community structure (e.g., Norby et al., 2019). C release via microbial respiration can be impacted by soil moisture (Blanc-Betes et al., 2016; Natali et al., 2015; Elberling et al., 2013), temperature (Hicks-Pries et al., 2013) and active layer depth (O’Donnell et al., 2011), as well as shifts in the quantity and quality of available organic matter (Treat et al., 2014; Hough et al., *in press*). Primary producers initially fix C and supply that C to the subsurface where it can be reworked by subsurface microorganisms. As the ultimate source of organic inputs to the peat, plants exert a strong control on SOM quantity and quality (Sutton-Grier and Megonigal 2011) which we hypothesize controls GHG production rates and their variation across thaw habitat types. Connecting the quality of aboveground C sources to differences in peat SOM is an essential step in testing the hypothesis that climate-associated vegetation shifts in peatlands influence changing C gas emissions.

Four major vegetation types dominate in high-latitude peatlands: bryophytes (mosses), graminoids (sedges), shrubs, and trees (Clymo 1987; Rodwell 1991). While warmer temperatures accelerate C loss from peat (Hopple et al., 2020; Hanson et al., 2020), this loss is greater when graminoids and shrubs dominate rather than *Sphagnum* mosses (Walker et al., 2016). *Sphagnum* is thought to suppress decomposition rates and thus GHG production by supplying poor-quality SOM (van Breeman 1995; Turetsky 2003), by microbial inhibition via acidification of the environment (Spearing 1972), and by production of inhibitory phenolic compounds (Rudolph and Samland 1985; Williams et al, 1998) and antimicrobial acids and sugar derivatives (Fudyma et al., 2019). Thus, environmental changes causing *Sphagnum* declines and increasing dominance by shrubs or sedges (e.g. McPartland et al., 2020; Norby et al., 2019; Walker et al., 2016; Johannson et al., 2006) is likely to result in more reactive and bioavailable SOM (Chanton et al., 2008; Tfaily et al., 2013; Wilson et al., 2021a). However, evidence has also been presented that compounds associated with some shrubs inhibit SOM degradation (Wang et al., 2021; 2015). Sedges, such as *Carex* and *Eriophorum*, have been correlated with higher CH_4_ production (Hines et al., 2008) and greater SOM reactivity (Chanton et al., 2008), thought to occur because sedges contain more cellulose, bioavailable N, and a higher proportion of labile compounds compared to *Sphagnum* (AminiTabrizi et al., 2020; Hodgkins et al., 2014, 2016). Graminoids also contain aerenchyma which are capable of transporting O_2_ to the rhizosphere, potentially enhancing decomposition. In contrast, *Sphagnum* lack such tissues, thus *Sphagnum*-dominated habitats generally have lower O_2_ availability providing a further thermodynamic constraint on SOM degradation in *Sphagnum*-dominated habitats.

Here, we investigate how permafrost thaw-driven changes in the quality of plant-derived organic matter influence SOM properties and thereby microbial decomposition. In this study, we analyze fresh plant material and peat collected from three habitat types across a thawing permafrost mire using the complementary techniques of Fourier Transform Infrared Spectroscopy (FT-IR) of solid phase material and Fourier Transform Ion Cyclotron Resonance Mass Spectrometry (FTICR-MS) of water extracts. These analyses will be used to assess (1) the quality of plant organic matter inputs in each of the three habitats, (2) which plant compounds accumulate as peat in each habitat type, and (3) the pathways by which plant-derived compounds are decomposed and how these differ across habitat types. Our assessments of organic matter quality will be used to determine how different plant types contribute to changes in SOM quality and drive GHG production rates across the thaw gradient. This information could be used to infer peatland-atmosphere feedback resulting from climate-driven shifts in plant community composition.

## Methods

### Site Description

Stordalen Mire (68.35°N, 19.05°E) is located in northern Sweden just north of the Arctic circle within the region of discontinuous permafrost. Climate change has accelerated thawing in the recent few decades leading to changes in hydrology and vegetation cover which have resulted in a patterned mosaic of habitat types within the mire (Johansson et al., 2006; Kokfelt et al., 2009); we focus here on the three dominant habitat types at the site: palsas, bogs, and fens. Some areas of the mire are still underlain by intact permafrost and elevated above the surroundings into relatively dry palsa plateaus. Warming has caused thawing of the permafrost in some areas causing, e.g., palsas to collapse and flood, producing wetter collapse features (Johansson et al., 2006). *Sphagnum* can infiltrate such pools, eventually elevating the surface enough to form a bog, or in some cases, the insulating effects of the *Sphagnum* are sufficient to allow the permafrost to refreeze. Alternatively, palsa can thaw completely and subside to the level of the surrounding water table, causing flooding and creating a fully-inundated fen. Fens are characterized by sedges and other aquatic vegetation (Zoltai 1993; Vitt et al., 1994; Jorgenson et al., 2001; Malmer et al., 2005), high CO_2_ uptake, and the highest CH_4_ emissions of the three habitat types (Hodgkins et al., 2014; McCalley et al., 2014). A bog, dominated by *Sphagnum*, can develop if the thawing permafrost collapses but remains above the local water table.

In addition to the hydrological differences, plant communities also change across this gradient of habitat types, from tundra-type vegetation dominated by shrubs, mosses, lichens, and small sedges in the dry palsa; to *Sphagnum* and small sedges in the bog; to tall sedges with some *Sphagnum* in the fen (Malmer et al., 2005). These differing plant communities likely contribute to differing SOM quality (Chanton et al., 2008; AminiTabrizi et al., 2020; Hodgkins et al., 2014, 2016; Tfaily et al., 2013), leading to much higher overall CH_4_ and CO_2_ emission rates from fens as compared to bogs (Hodgkins et al., 2014) and the even-drier palsas (McCalley et al., 2014). Since the 1970’s, the areal coverage of *Sphagnum* across the mire has declined significantly (Malmer et al., 2005), giving way to increased sedge cover as wetter conditions across the mire have increased the areal coverage of fen habitats (Kokfelt et al., 2009; Bäckstrand et al., 2010). This gradient in habitats across the mire creates a unique opportunity to explore changes in SOM quality with habitat transition within the context of changing greenhouse gas production rates.

### Plant Collection

To explore differences in plant organic matter inputs across the three habitat types, samples of the characteristic species from each habitat (Malmer 2005) were collected. Water extracts from the whole plants and tissue types (leaves, stems, roots) were used to compare organic matter inputs composition across the different plant types. Plant-associated compounds were then compared to the peat from each habitat to understand what compounds were easily decomposed (i.e., which compounds stimulated microbial activity) versus those compounds that were less bioavailable and that tended to accumulate in the peat. Plants were collected during the peak of the growing season (early August) in 2014 resulting in the following samples for each habitat: palsa – *Rubus chamaemorus, Betula nana, Empetrum nigrum, Andromeda polifolia, Dicranum elongatum, Eriophorum vaginatum*, fruticose lichen of unknown species; bog – *Sphagnum spp*.; fen – *Eriophorum angustifolium, Carex rostrata*. Whole plant samples were collected and separated by tissue type (roots, stems, and leaves), then immediately flash-frozen in liquid N_2_ and kept frozen at -20°C until processing in February 2015. Since mosses do not have root, stem, and leaf differentiation, they were not separated and were processed as whole plants. Additional plant samples for FT-IR analysis were collected in August 2015 and included *Sphagnum fuscum, S. magellanicum, E. nigrum, A. polifolia*, and an unknown species of lichen. These samples were similarly flash frozen in the field in liquid N_2_ and then kept at -20°C until analysis.

### Peat Collection

Peat was collected in August 2014, from the same three habitats along the thaw gradient where plants were collected, using a Wardenaar corer (Eijkelkamp, Raleigh, NC USA). The cores were sectioned by depth, placed in Teflon coated vials, and frozen at -20°C before analysis. On returning to the lab, peat samples were freeze dried and ground to a homogenous powder using a SPEX SamplePrep 5100 Mixer/Mill ball grinder. Porewater was also collected from the site using a perforated stainless-steel tube inserted into the peat to the desired depth. Gentle suction was applied using a gas tight syringe fitted to the tube using a three-way valve. Once 30 mL of porewater was obtained, it was placed in a polycarbonate sample vial and frozen at -20°C prior to analysis. An additional 30 mL of porewater was collected in three locations within 1 m of the core for replicate pH analysis immediately in the field. Samples were collected from the shallowest depth it was possible to draw porewater: 10-14 cm in the bog and 1-5 cm in the fen. We used the solid peat to compare the compounds present in the palsa, where the conditions are not water saturated and no porewater could be collected, to the other sites where water saturation has already effectively extracted dissolved compounds from the peat. A list of sample types and the number of samples analyzed by each method are given in Supplemental Table 1.

### Fourier Transform Infrared Spectroscopy (FT-IR)

To examine the bulk chemical characteristics of the plants and solid peat, the dried and ground material were analyzed by Fourier Transform Infrared Spectroscopy (FT-IR). For FT-IR, only stems and leaves from each plant were available for analysis (no roots). Recent advances in FT-IR analysis allow us to quantitatively evaluate differences in carbohydrates and aromatic compounds among samples (Hodgkins et al., 2018). FTIR spectra were collected using a PerkinElmer Spectrum 100 FTIR spectrometer fitted with a CsI beam splitter and a deuterated triglycine sulfate detector. Transmission-like spectra were obtained using a Universal ATR accessory with a zinc selenide/diamond composite single-reflectance system. Each sample was placed directly on the ATR crystal, and force was applied so that the sample came into good contact with the crystal. Spectra were acquired in % transmittance mode between 4000 and 650 cm^−1^ (wavenumber) at a resolution of 4 cm^−1^, and four scans were averaged for each spectrum. The standard deviations of carbohydrate and aromatic carbon values were within 5% of the mean values when 4 replicate samples were run and scanned four times. That is, if a sample was found to be 30% carbohydrate, the analytical error on 4 aliquots that were each scanned 4 times was 1.5%. Spectra were ATR-corrected, baseline-corrected, and then converted to absorbance mode using the instrument software. Area-normalized and baseline-corrected peak heights for common classes of compounds observed in SOM were calculated using the methods and script described by Hodgkins et al., (2018), expanded to include peak assignments by Palozzi and Lindo (2017).

### Fourier Transform Ion Cyclotron Resonance Mass Spectrometry (FTICR-MS)

We used Fourier Transform Ion Cyclotron Resonance Mass Spectrometry (FTICR-MS) to gain a higher resolution view of the compounds present in the palsa peat, peat porewater from the bog and fen, and the plant samples. Plant samples were thawed and each tissue type (roots, stems, and leaves when available), in addition to whole plant samples for mosses, which lack leaf/stem/root differentiation, were analyzed after water extraction in which 0.5 g of undried plant material was shaken in 4 mL nanopure water and then allowed to sit for 2 hours, and the supernatant decanted. The resulting extracts were mixed 1:2 with HPLC-grade methanol and immediately direct-injected into a 12 T Bruker ESI-FTICR-MS spectrometer operating in negative mode. Solid peat samples (from an 8-18 cm deep section at each site) were analyzed after water-extracting the dried and ground peat samples. For this method, 0.5 g of the dried and ground peat, which is expected to yield 25 mg C, was added to 1 mL of degassed deionized water and then placed on a shaker for 2 hours. The solutions were then centrifuged to form a pellet and the supernatant was decanted. The supernatant and porewater samples were then each mixed 1:2 (by volume) with HPLC grade methanol, and the resulting solutions were injected through direct injection onto a 12 T Bruker ESI-FTICR-MS spectrometer operating in negative mode. For each sample, ninety-six individual scans were averaged and then internally calibrated using organic matter homologous series separated by 14 Da (i.e., CH_2_ groups). The mass measurement accuracy was <1 ppm for singly charged ions across a broad m/z range (i.e., 200 < m/z < 1200). Chemical formula assignments were made using an in-house built software program following the Compound Identification Algorithm, described by Kujawinski and Behn (2006) and modified by Minor et al., (2008) and based on the following ‘Golden Rules’ criteria: signal/noise > 7, and mass measurement error < 1 ppm, taking into consideration the presence of C, H, O, N, S and P and excluding other elements. All observed ions in the spectra were singly charged based on identification of 1.0034 Da spacing found between carbon isotopologues of the same molecule (e.g., between ^12^C_n_ and ^12^C_n-1_– ^13^C_1_). Two technical replicates were collected for most samples and, when available, peaks present in either (or both) spectra were combined and the signal intensities were averaged for downstream analysis.

Complex organic matter such as both the plant extracts and the peat are expected to result in thousands, if not tens of thousands, of unique compounds by FTICR-MS. A number of approaches exist to aid in visualizing such complex datasets. These include the use of van Krevelen diagrams that depict the H/C vs. O/C ratios of individual compounds, which enables tentative inferences about general compounds classes. For example, lipids are generally low O/C with high H/C, while carbohydrates generally fall in the region near O/C = 1 and H/C = 2. In addition, the molecular formulae derived from FTICR-MS analyses can be used to calculate the nominal oxidation state of the carbon (NOSC) in individual compounds observed in the DOM. This is done through a simple calculation from the molecular formula NOSC = 4 – (4C + H – 3N – 2O + 5P – 2S)/C (Keiluweit et al., 2016), but provides tremendous insight into the thermodynamic energy yield on oxidation of that C (La Rowe and van Cappellin 2011), which is directly relevant to understanding organic matter quality (Wilson and Tfaily 2018).

### Chemical transformation Analysis

Chemical transformation analysis of the chemical compounds identified by FTICR-MS involves calculating the mass differences between individual compounds and matching those mass differences to specific chemical moieties. By matching these results with known biochemical transformations accomplished by microorganisms in the environment, it is possible to infer the decomposition pathways by which individual compounds are degraded and produced (e.g., Stenson et al., 2003; Kujawinski et al., 2016; Wilson et al., 2017). This process is possible because of the extremely high mass resolution of the FTICR-MS technique which allows us to narrow down the possible matches within 1 ppm. The current database of microbial transforms contains 186 unique transforms (Wilson et al., 2017), including hydroxylation, methoxylation, and transamination reactions.

## Results

The pH for the porewater at the bog surface averaged 4.2 ± 0.2. In the fen, the porewater pH at the surface averaged 5.6 ± 0.4. No porewater was available in the surface palsa for collection.

### FT-IR

The leaf and stem FT-IR spectra were quite similar for both vascular plants (*E. nigrum, A. polifolia*) in the palsa habitat, with the exception that *A. polifolia* leaves had lower carbohydrate content and *E. nigrum* leaves had lower carboxylic acid content and aliphatic waxes compared to stems from the same plant (Figure 1; Supplemental Table 2). Because of the similarity between leaf and stem spectra for each plant and because of expected higher turnover of leaves compared with stems, we compared the FT-IR spectra from the leaves of the dominant plants to the peat in each habitat type (Figure 1). High peak intensities were observed at wavenumbers corresponding to carbohydrates (i.e., O-alkyls at 1030 cm^-1^); lignin (1265 cm^-1^); humic acids (1426 cm^-1^); phenolic lignin-like structures (1515 cm^-1^); protein-like (1550 cm^-1^); aromatics (1650 cm^-1^); C=O stretching associated with carboxylic acids, aldehydes ketones and other oxygenated moieties (1720 cm^-1^); and aliphatic fats (2920 cm^-1^ and 2850 cm^-1^)(Supplemental Table 2). In the palsa, the two depths of peat were very similar, with deeper peat having a slightly higher aromatic, carboxylic acid, and waxy lipid content relative to the shallower peat (Figure 1d). The carbohydrate content of the palsa peat was lower than the S. fuscum and lichens. The waxy lipids, peaks 2850 cm^-1^ and 2920 cm^-1^, were extremely well differentiated in the leaves of *E. nigrum* and *A. polifolia* compared to the leaves of *S. fuscum* and lichens (Figure 1a).

**Figure 1:**
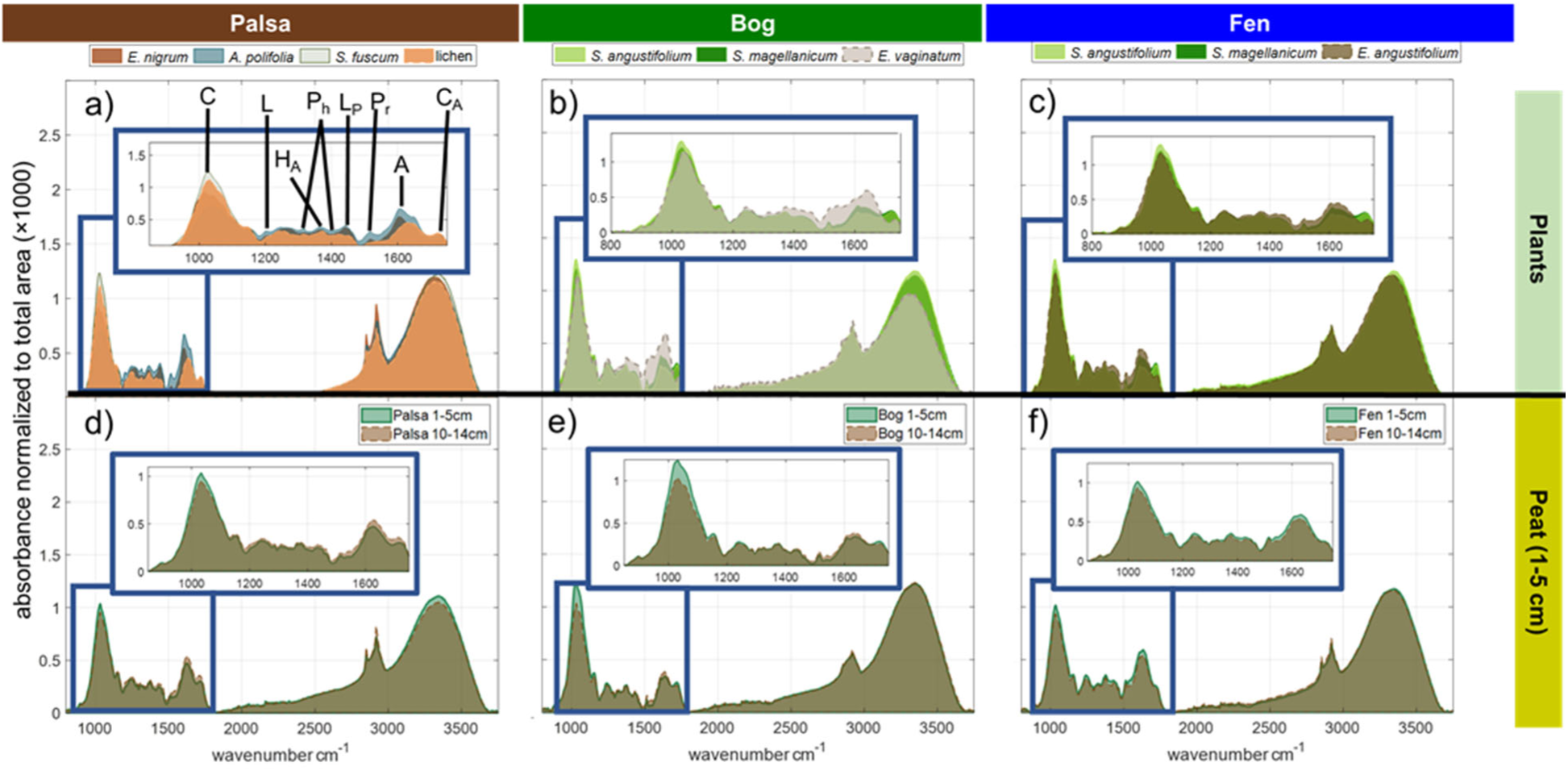
Layered FT-IR spectra comparing dominant plants (leaf material only) and peat in each habitat. All spectra are baseline-corrected and normalized to total peak area as described in Hodgkins et al., (2018). Inset plots enlarge the 850-1750 cm^-1^ region where many chemical functional groups exist within a short span of wavenumbers. In the panel (a) inset, important peaks discussed in the text are marked: C = carbohydrates, L = lignin, H_A_ = humic acids, P_h_ = phenolic-lignin, L_P_ = lignin-like, P_r_ = proteinaceous, A = aromatics, C_A_ = carboxylic acids. Panel (a) shows the overlaid spectra for lichens, A. *polifolia, E. nigrum, and S. fuscum*, the dominant plant types in the palsa, while palsa peat spectra are shown in panel (d). Panel (b) shows the overlaid spectra for the bog’s dominant plants *S. angustifolium, S. magellanicum*, and *E. vaginatum*, with the bog peat in panel (e). Panel (c) shows the fen’s dominant plants, *E. angustifolium, S. angustifolium, and S. magellanicum*, which are reflected in fen peat in panel (f).

In the bog, *Sphagnum* mosses and *E. vaginatum* leaf and bog peat FT-IR spectra were compared (Figure 1b,e). The waxy lipid peaks at 2850 and 2920 cm^-1^ were slightly more differentiated in the *E. vaginatum* compared to the *Sphagnum*, consistent with higher waxy lipid content in *E. vaginatum*. The leaf carboxylic acid peak (1720 cm^-1^) was stronger in the *Sphagnum* compared to the *E. vaginatum*. The bog peat had the highest humic acid content of any of the sites (Supplemental Table 2).

In the palsa, *E. vaginatum* had the highest leaf aromatic content (Supplemental Table 2). In the FT-IR spectra of the peat from the different habitats (Figure 1d, e, f) several absorption bands typical of humic materials were observed in our samples (Artz et al., 2008; Chapman et al., 2001; Leifeld et al., 2012). The bog peat had the highest carbohydrate content of any of the sites. The fen peat had a higher abundance of protein-like structures and lower abundance of carboxylic acids compared to the other sites. Aliphatics (2920 cm^-1^ and 2850 cm^-1^) were much less well defined in the bog peat compared to the other sites, indicating fewer waxy lipids (Artz et al. 2008; Cocozza et al. 2003) compared to the other sites. Comparisons between the shallow and deep peat (Figure 1d,e,f) at each site indicated differences in decomposition among the habitats. Carbohydrates, protein-like structures, and aromatics became less pronounced with depth in the fen, while aliphatic waxes increased and humic acids decreased with depth in the bog.

### FTICR-MS Results

Among all of the plant samples, leaves, stems, and roots combined we observed 19,072 compounds via FTICR-MS. Of those, we were able to assign a molecular formula to 14,260 compounds (75%), which is a typical assignment rate for complex SOM. Across all habitats in the peat, we observed 15,198 unique compounds of which we were able to assign molecular formulae to 11,254 (74%). Palsa plants had the highest diversity of compounds (n = 11,633, Figure 2). Of those, the majority of compounds were not present in the peat (75%) suggesting that they were microbially decomposed and/or processed. The remaining 25% were present in the peat, suggesting that they are resistant to microbial decay and accumulate over time (Figure 2). The bog plants had the lowest diversity of compounds (n = 3578) but they also appeared to be the most resistant to microbial decomposition as 47% were observed in the bog peat. The fen was intermediate between the palsa and bog with fen plants having 5312 compounds of which 41% were observed to accumulate in the peat (Figure 2).

**Figure 2:**
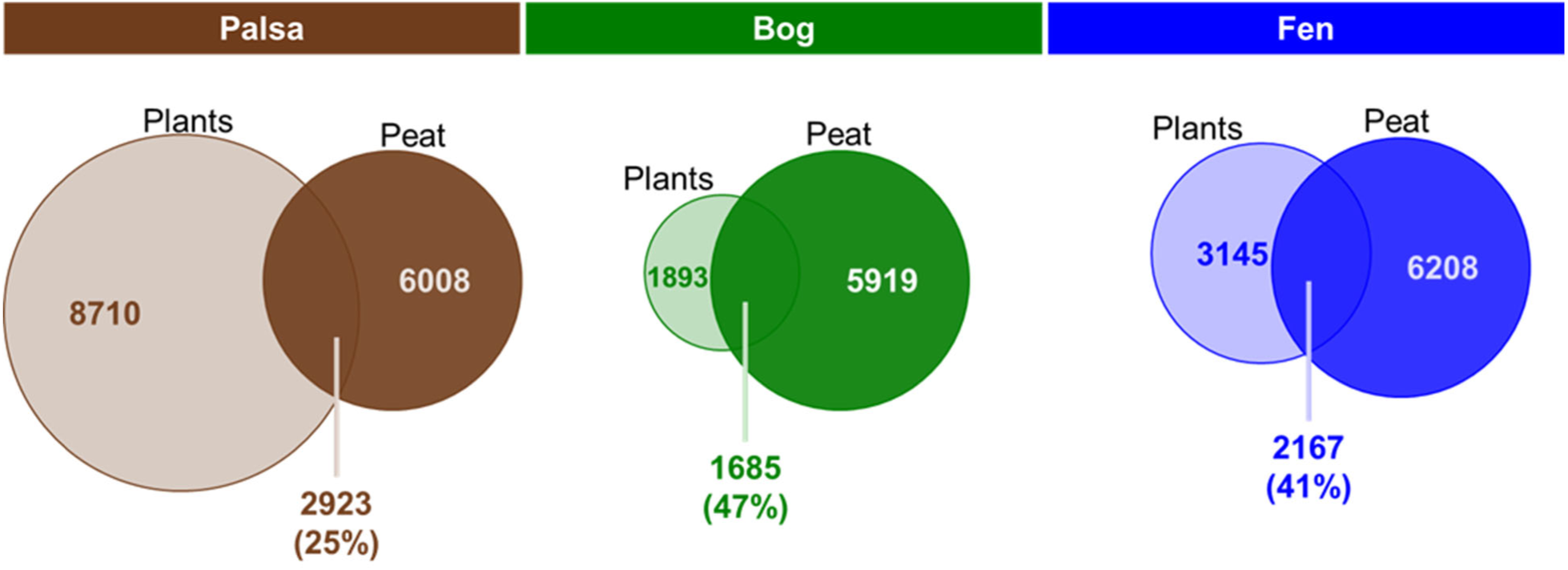
Comparison of compounds in plant extracts (leaf, roots, stems, and whole mosses combined) and in the shallow peat. Numbers in each circle indicate the number of different compounds identified by FTICR-MS that are unique to either the plants or peat collected from each habitat, while shared compounds are indicated by the overlapping regions (with numbers directly below, and the percentage of plant compounds these represent). We refer to these overlap-region compounds as “accumulated” because they are plant-derived and appear resistant to microbial decomposition, persisting in the peat.

We compared the compounds observed in the composite plant extracts (leaves only, or whole plants for lichens and mosses) to those in the shallow peat from each habitat type (Figure 3) (The supplement contains comparisons of the stem extracts from the palsa, Supp. Fig. 1, and root extracts to the peat for each habitat, Supp. Fig. 2). There were striking differences in the plant leaf compounds as well as the peat across the different habitats (Figure 3). The plant composite in the fen (including extracts of *E. angustifolium*, and *C. rostrata*) had low numbers of lipid-like (fatty acids) and protein-like compounds relative to the dominant plants from the palsa (lichens, *A. polifolia*, and *E. nigrum*) and bog (*Sphagnum* and *E. vaginatum*) and habitats (Figure 3). Both the bog and fen peat had relatively more tannin-like compounds compared to the palsa peat despite having lower abundances of these compounds in their plant composites relative to the palsa.

**Figure 3:**
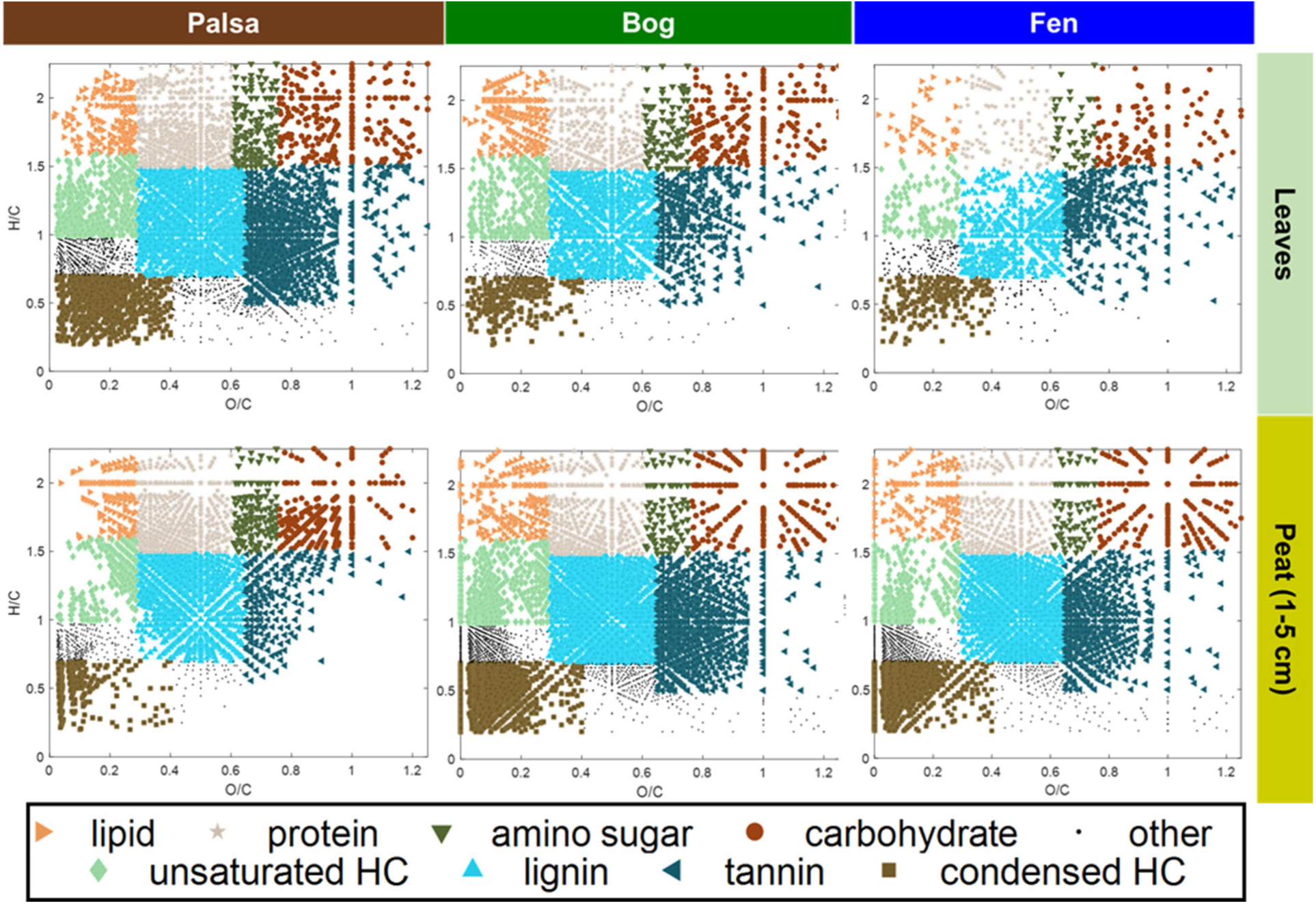
The combined FTICR-MS of the dominant plant species’ leaves (or for mosses and lichens, the entire plant) and near-surface (1-5 cm) peat extracts from each habitat. Dominant plants for the palsa were lichens, *A. polifolia, and E. nigrum*; for the bog *Sphagnum* and *E. vaginatum*; and for the fen *E. angustifolium* and *C. rostrata*. Each point represents an observed molecular formula, and is categorized by color and symbol according to inferred compound classes.

We calculated the nominal oxidation state of the carbon (NOSC) in the water extracts pf the dominant plant leaves (whole plants for lichens, *Sphagnum*) from each habitat (Figure 4) as a metric for determining organic matter quality (Wilson and Tfaily 2018). Lichens, *A. polifolia*, and *E. nigrum* together comprise 31% of the aboveground leaf, 95% of aboveground stem, and 22% of the belowground (root) biomass in the palsa. *Sphagnum* accounts for 74% of the biomass in the bog overall, with *E. vaginatum* contributing 13% of the bog’s aboveground and 20% of the belowground biomass. In the fen, *E. angustifolium* is 63% of the aboveground and 81% of the belowground biomass, while *C. rostrata* contributes approximately 5% of the above and belowground biomass. Sphagnum had the lowest NOSC of any of the habitat-dominant plants (Figure 4). *E. angustifolium*, in the fen, had intermediate NOSC values that were nevertheless significantly higher than those found in *Sphagnum*. The palsa plant community was more diverse, lichens had the highest NOSC values and *E. nigrum* and *A. polifolia* had significantly higher NOSC values than *Sphagnum*.

**Figure 4:**
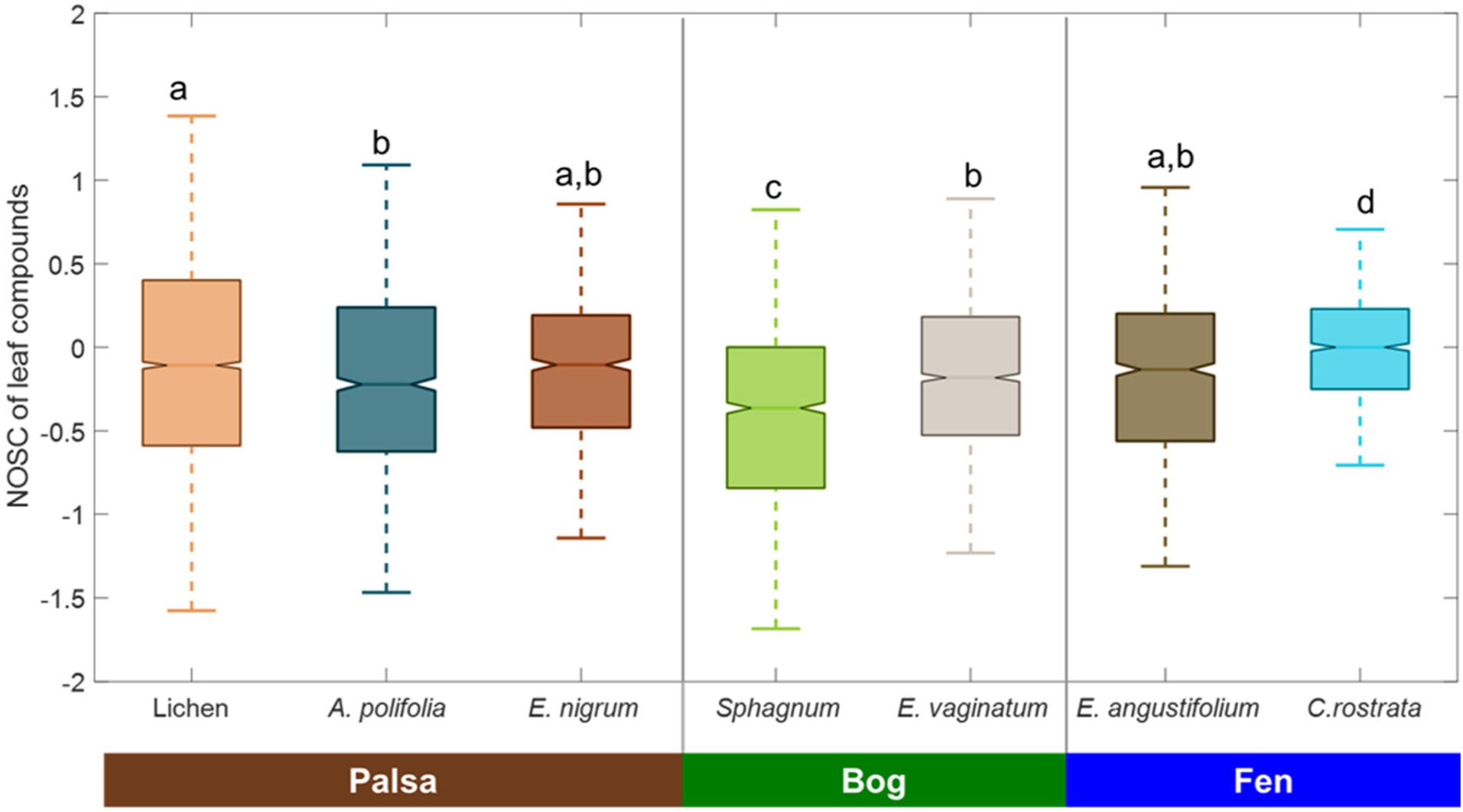
Nominal oxidation state of carbon (NOSC) for dominant plant leaf extracts (whole plants for lichens, *Sphagnum*) from each habitat. Different lowercase letters above bars indicate significant differences by ANOVA followed by pairwise comparison (Tukey’s Honestly Significant Difference).

It is possible that the average NOSC was being disproportionately influenced by a large number of compounds with extreme NOSC values, but that were present at overall low concentration. To determine whether this was the case, we calculated the normalized signal intensity for each compound in the composite of all plant parts (leaves, stems, and roots combined) from all plants collected in each habitat. We plotted the cumulative normalized signal intensity against the NOSC of the compounds (Figure 5) and found that compounds with NOSC < 0 accounted for 46% of the signal intensity in the palsa, 71% of the signal intensity in the bog, and 58% of the signal intensity in the fen. Although not strictly quantitative within similar sample types, signal intensity roughly follows concentration in samples with similar overall matrices.

**Figure 5:**
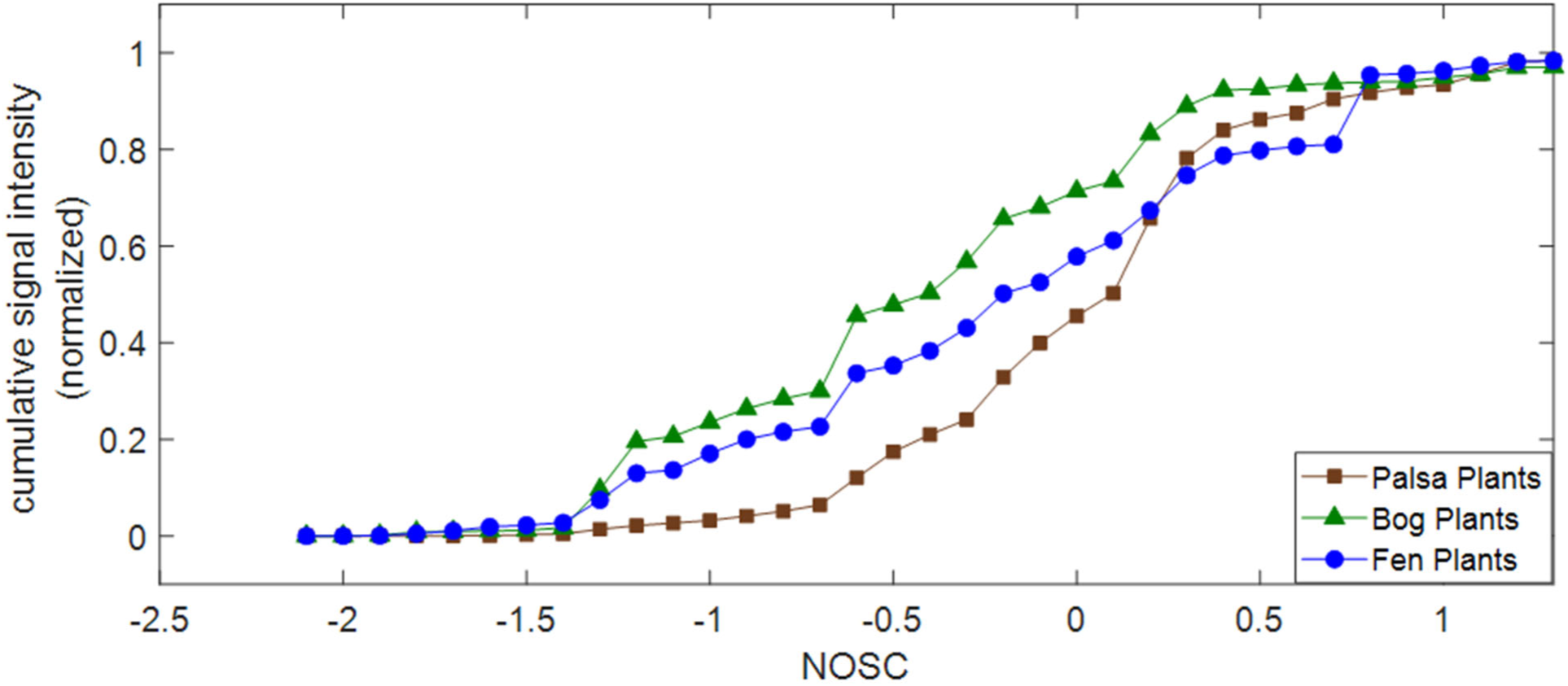
Cumulative signal intensity normalized to total intensity as a function of NOSC in the composite plant samples from each habitat.

We then examined the characteristics of the plant-associated compounds that were either (1) consumed or that (2) accumulated in the peat as well as the compounds in the peat that were not present in the original plant material and were therefore assumed to be (3) microbially produced (Figure 6). Consumed compounds are those that appear in the plant extracts but not in the peat suggesting that they were decomposed. Accumulated compounds appear in both the plants and the peat, apparently resistant to decay. Some compounds appear only in the peat suggesting that they were produced during decomposition presumably by microbial processes, although abiotic production is also possible (Fudyma et al., 2020). This comparison is sensitive to even minor abundance plant compounds, for example, compounds could appear to be produced if they came from a minor species that was not included in the plant mixture. To minimize this effect, we included all plant parts (leaves, stems, and roots) from all plant species sampled at a given habitat (regardless of abundance) to compare against the peat compounds. In the palsa, this included: lichens, *A polifolia, E. nigrum, D. elongatum, R. chamaemorus*, and *B. nana*. In the bog this included *Sphagnum, E. vaginatum*, and *E. angustifolium*. In the fen this included *E. angustifolium, C. rostrata*, and *Sphagnum*. In the palsa, tannins, lignins, condensed hydrocarbons, and carbohydrates were more readily consumed compared to the bog and fen. In contrast, a higher percentage of the total lipid content was consumed in the bog and fen compared to the palsa. In the palsa a lower percentage of the lignins, tannins, and condensed hydrocarbons accumulated in the peat (Figure 6) suggesting that the higher redox in the palsa facilitated the decomposition of these types of compounds. Compared to palsa, a high proportion of condensed hydrocarbons, tannins, and lignins appeared in the bog and fen (Figure 6 c) peat. In addition to unclassified (‘other’) compounds, a high proportion of protein-like compounds were produced in the palsa compared to the other sites (Figure 6 c).

**Figure 6:**
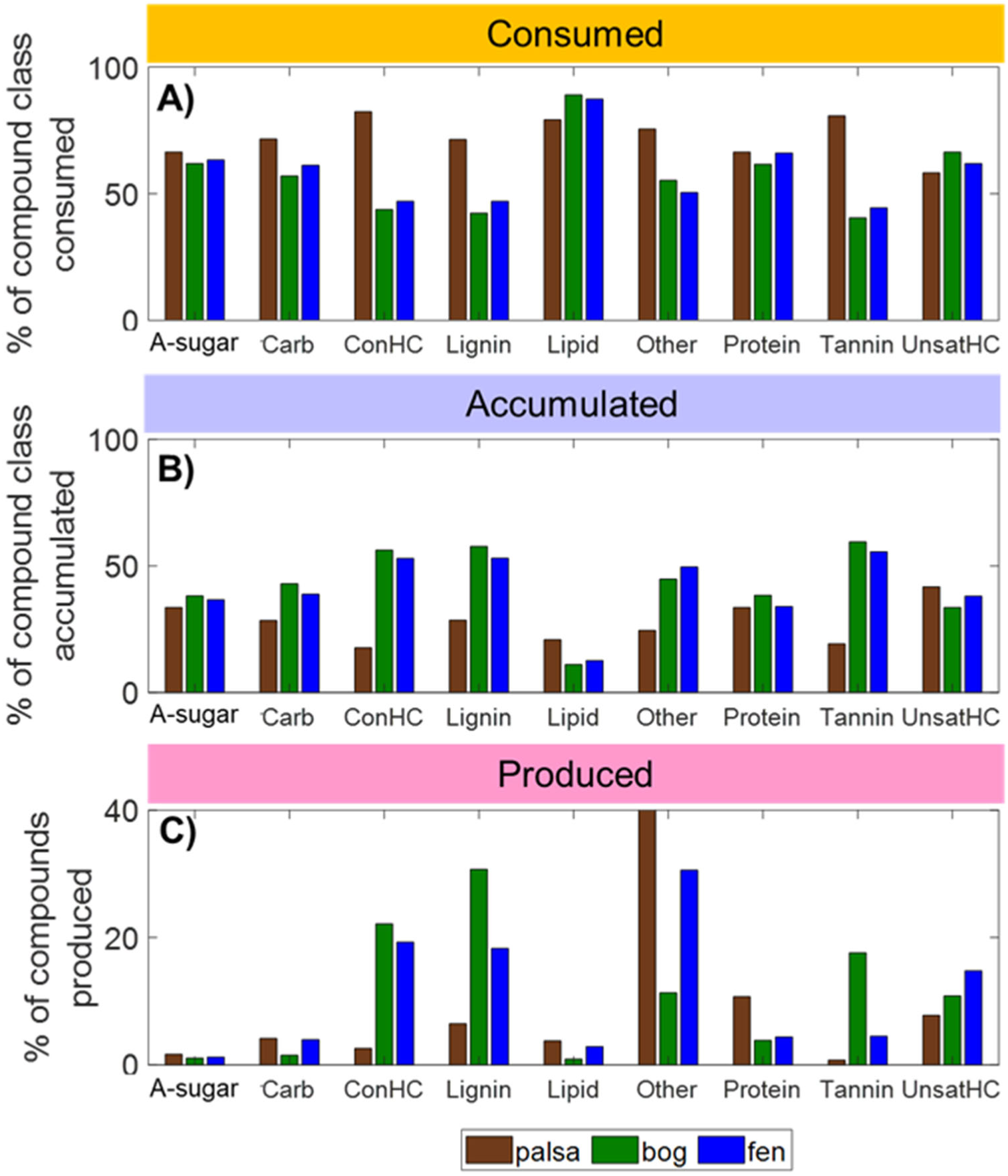
Inferred percentages of consumed, accumulated, and produced compounds within the peat extracts, by chemical class (inferred from the molecular formulae from FT-ICR MS, per Figure 3), for each habitat. (A) Percentage of consumed compounds, calculated as the number of consumed compounds in each class divided by the total number of compounds in that class originally present in the plant extracts. (B) Percentage of accumulated compounds, calculated as the number of plant compounds of each class that were also present in the peat, divided by the total number of compounds in that class originally present in the plant extracts. (C) Percentage of produced compounds, calculated as compounds present in the peat but absent from the plant material, and are inferred to be microbially produced, divided by the total number of compounds in the peat. Note that the maximum y-axis value in (C) was set to 40% to improve visualization of differences; the only produced class that exceeded 40% was the unclassified “other” compounds at 62.5% in the palsa.

We also examined the nitrogen (N), sulfur (S), and phosphorus (P) content of the various compounds (Figure 7). Overall, the plants in the fen had a higher proportion of both nitrogen and sulfur containing compounds compared to the plants in either the bog or palsa. In the bog, a higher percentage of nitrogen-containing compounds was consumed relative to either the fen or the palsa (Figure 7). Additionally, when compared to the proportions present originally in the plants, a higher percentage of N- and S-containing compounds were consumed in the bog. A higher percentage of CHO-only compounds accumulated in the fen compared to the starting plant material suggesting that N-, S-, and P-containing compounds were preferentially consumed. A high percentage of the produced compounds in the palsa were CHO-only compounds and very few phosphorus or sulfur containing compounds were produced. This result contrasts with the bog and fen where phosphorus and sulfur containing compounds were ∼40% of the total compounds produced.

**Figure 7:**
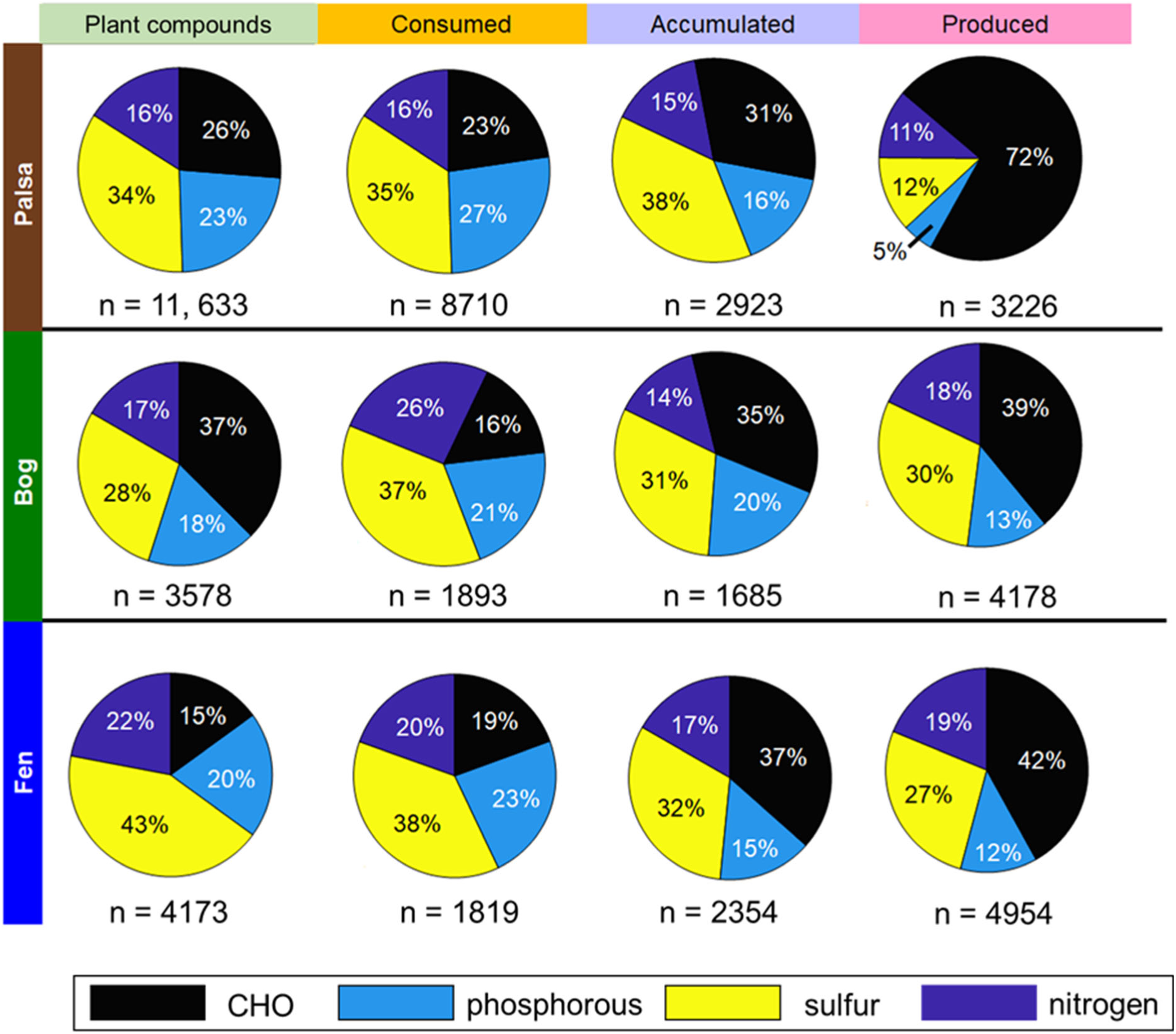
The proportion of compounds that were exclusively CHO or that contained N, S, or P, in each habitat, for total plant compounds (for leaves, stems, and roots of the plants collected in each habitat), and those inferred to have been consumed, accumulated, or produced in the peat. Percentages the number of compounds in each category, divided by the total number of compounds in the group (n) indicated under each chart.

To understand potential differences in the decomposition pathways among the three habitats that have contributed to the differences observed in the produced compounds, we calculated the number of times each transform (i.e., chemical transformation pathways by which SOM decomposes) occurred within a sample in the peat and plotted the most frequently observed transforms from each site (Figure 8). Hydrogenation (H_2_) was the most frequent transform for all of the habitat types. Demethylation followed by oxidation (CH_2_-O) was the second most frequent for the palsa and fen, but side-chain (de)methylation (CH_2_) was second for the bog. Transformations involving changes of N (OH-N, O-NH, and NH_3_-O) were highest in the palsa.

**Figure 8:**
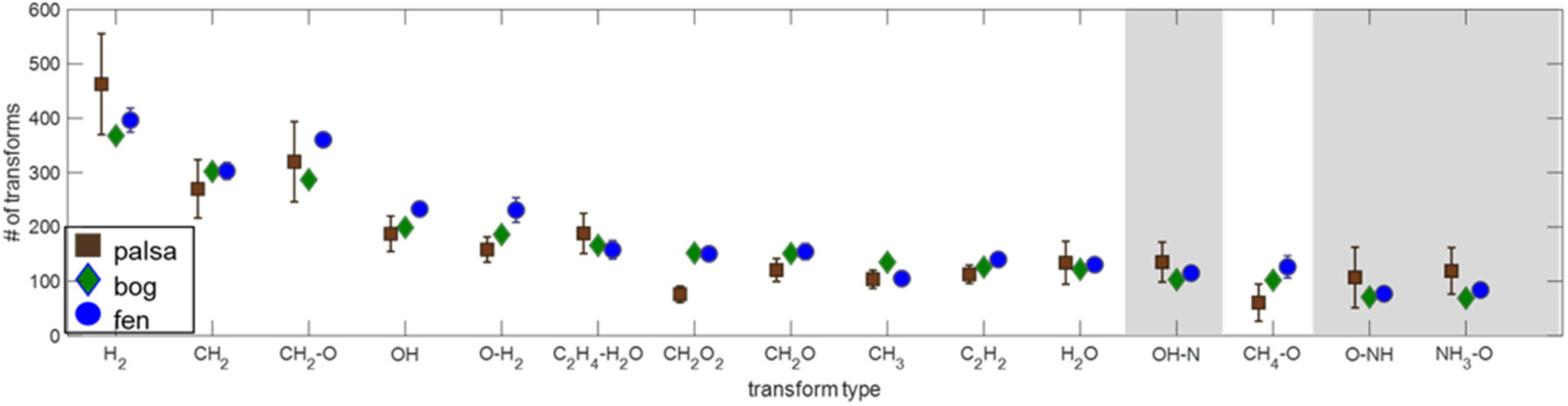
Top transforms for each habitat’s peat given as the molecular differences among compounds (i.e., H_2_ refers to a difference of 2 hydrogen atoms). Points are plotted as averages ± 1 s.d. for 3 samples of palsa, and for 2 samples of fen; one sample was available for bog. Transforms involving nitrogen are highlighted in gray.

## Discussion

In our investigation of changes in plant and soil organic material (SOM) composition along a permafrost thaw gradient, we observed a strong relationship between the plant-derived organic compounds and SOM compounds suggesting that aboveground vegetation and roots shape belowground processes and subsequent SOM decomposition in this peatland. Across the thaw gradient, there were significant changes in CO_2_ and CH_4_ production potential (Hodgkins et al., 2014) and emissions (McCalley et al., 2014). The palsa was associated with net CO_2_ emission and little, or no, CH_4_ production, the bog and fen both exhibit net CO_2_ uptake, and CH_4_ emissions from the fen were the highest of the three habitat types (Bäckstrand et al., 2010; McCalley et al., 2012). CO_2_:CH_4_ ratio production potentials, clearly indicated that the fen was the most methanogenic of the three sites (Hodgkins et al., 2014).

Hodgkins et al. (2014) found that differences in the major SOM classes drove variability in greenhouse gas (GHG) emissions across the mire. In particular, they ascribed increasing GHG emissions across the thaw gradient to increasing SOM lability as inferred from decreasing C/N ratios and lower molecular weight, aromaticity, organic acid, and organic oxygen contents. Our results support that conclusion and further show that differences in SOM are driven by changes in the plant community, confirming the central role of plants in controlling SOM quality. The plant community is the initial source of the organic matter to the subsurface (Sutton-Grier and Megonigal, 2011). Microbial decomposition then removes some chemical species while creating others, thereby modifying the inputs in a way that is partially dependent on oxygen availability within each habitat. *Sphagnum* plays a particularly strong role in habitats where this species dominates. Organic matter in *Sphagnum* extracts has significantly lower NOSC than other abundant plant species (Figure 4). Such low NOSC is consistent with low organic matter quality (Wilson and Tfaily 2018) suggesting a mechanism for suppressed SOM decomposition in the bog, especially as compared to the fen. In addition, *Sphagnum* produces many compounds that are potentially inhibitory to microbial activity (Fudyma et al. 2020) including organic acids which result in lower pH in the bog relative to the fen habitat. All of these factors work synergistically to facilitate C storage in *Sphagnum*-dominated environments. The percentage of plant compounds that accumulate in the peat, and are therefore less bioavailable, increases from palsa (25%) to fen (41%) to bog (47%) (Figure 2), which is opposite to the trend in plant species diversity across the habitats. *Sphagnum* limits decomposition rates by producing low NOSC compounds (Figure 4), producing microbially inhibitory compounds (Fudyma et al., 2020), but it appears that these effects of *Sphagnum* can be attenuated by increasing proportional cover of other plant species.

The high carbohydrate peak in the FT-IR analysis of the bog peat relative to the other habitats (Figure 1) was unexpected, particularly since CO_2_ production potential is low in the bog (Hodgkins et al., 2014) and carbohydrates are a highly bioavailable C source that should stimulate production. However, the high carbohydrate content of the bog peat is consistent with other observations that sugars tend to accumulate in *Sphagnum*-dominated peat (AminiTabrizi et al., 2020), and that the hydrolytic enzymes responsible for the initial breakdown of carbohydrates are less active in bog peat relative to the other habitat types (Woodcroft et al., 2018). Analysis of the solid phase (FT-IR) bog peat revealed high abundance of bioavailable carbohydrates, however, the FTICR-MS analysis revealed low quality organic matter in the water-soluble fraction. This implies that the availability of labile C in the bog is, in part, limited by solubilization of the cell walls, likely due to the low pH in the bog (pH = 4.2) which is known to inhibit DOM hydrolysis (Curtin et al., 2016). The high relative abundance of the carboxylic acid peak in the solid *Sphagnum* (Figure 1) is consistent with the high abundance of carboxylated sugars and uronic acids that comprise the structural components of Sphagnum cell walls (Painter 1991; Ballance et al., 2007) which could explain the relatively high carbohydrate peak in the solid bog peat as well as the lower pH in the bog relative to the fen.

In the fen, FT-IR results show a greater reduction in the intensities of nearly all of the important functionalities with depth of the peat compared to any of the other habitats (Figure 1f) which we conclude reflects higher decomposition rates. In both the palsa and fen it appears that decomposition rapidly follows organic matter deposition, based on the low abundance of carbohydrates even in the shallow peat relative to the available plants. In contrast, the bog peat intensities are quite similar to those of the *Sphagnum*, suggesting that the organic matter has changed (decomposed) little following deposition. This result is consistent with the lower GHG emissions in the bog relative to fen (McCalley et al., 2014; Hodgkins et al., 2014) and the supposition that bog SOM is less bioavailable than fen SOM.

We found a high abundance of waxy lipids in the leaves of *E. nigrum* and *A. polifolia* from the palsa (as seen in the strong differentiation between the 2850 cm^-1^ and 2920 cm^-1^ FT-IR peaks; Figure 1a) (Artz et al. 2008; Cocozza et al. 2003). While these compounds are frequently thought to be refractory, they do not appear as strongly even in the shallow peat, suggesting that they are at least partially degraded following deposition. Alternatively, because the leaves of *E. nigrum* and *A. polifolia* are very small and evergreen, they are likely underrepresented in the peat since they don’t all drop in the fall (unlike the deciduous plants in the habitat). The low differentiation between these peaks in the lichen is consistent with lichens lacking the waxy cuticle that coats plant leaves. The low relative abundance of carbohydrates in the palsa peat relative to the abundant plant species is consistent with fast decomposition in the surface depleting highly bioavailable compounds soon after deposition.

While FT-IR is practical for looking at overall changes of functional groups in the bulk solid-phase SOM, FTICR-MS provides finer-resolution detail of the water-extractable compounds, down to the individual molecular level. Overall, we observed fewer unique compounds in peat samples relative to the plant sample set as determined by FTICR-MS (15,198 vs 19,072 respectively). This result is consistent with loss of compounds with decomposition of the plant material following deposition. The percent of plant compounds that were also found in the peat increased from palsa (25%) to fen (41%) to bog (47%) (Figure 2). This pattern suggests that a higher percentage of plant compounds were decomposed in the palsa so that they are no longer detectable. The apparently higher decomposition in the palsa occurs even though the palsa also has the greatest number of different compounds of all the sites. The richness (i.e., number) of compounds observed in the plants across the different habitat types (Figure 2) follows the same pattern as the plant species diversity changes across the sites: palsa > fen > bog (Hough et al., 2020; Johansson et al., 2006). Interestingly, this trend is opposite that observed in the diversity of the plant-associated microbial communities across these sites (Hough et al., 2020; Wilson et al., 2021b).

Nevertheless, the richness of compounds in the peat is similar across the different habitats, which suggests that a high diversity of microbial pathways in the bog and fen is responsible for transforming the less diverse plant matter into more diverse peat. While there is considerable overlap in common compounds between the peat and the dominant plant types found within each habitat, many of the plant compounds were not found in the peat, while the peat also had many unique compounds not found in the plants. These results indicate both loss and production of novel compounds following plant organic matter deposition, presumably through the metabolic action of microorganisms. Only 25% of compounds from the palsa plant composite were also observed in the shallow peat (Figure 2), indicating that 75% of plant compounds were either consumed or metabolically processed into other molecules and that the plants in the palsa were largely bioavailable and susceptible to decomposition. It is likely that the higher lability (as inferred from NOSC) of the dominant plant compounds (Figure 4) contributes to the greater decomposition of organic matter from palsa plants. Additionally, the higher availability of oxygen as a terminal electron acceptor (TEA) in the palsa compared to the other sites could catalyze the decomposition of a range of bioavailable compounds in the palsa relative to the other habitats. The higher oxygen content could explain why hard to decompose chemical classes such as tannins, lignins, and condensed hydrocarbons, which accumulate in the bog and fen, are more readily consumed in the palsa (Figure 6).

In highly oxygenated environments, production of CO_2_ is thermodynamically favored, but in anoxic, TEA-depleted, waterlogged environments, CO_2_ is sometimes the only available TEA, resulting in CH_4_ production. Plants exert a strong influence on the CO_2_:CH_4_ ratio by being the prime source of organic substrates (i.e., electron donors) in the subsurface (Megonigal et al., 2004; Sutton-Grier and Megonigal 2011), and by controlling the availability of TEAs used in decomposing that organic matter. There is a strong relationship between NOSC calculated from the molecular formula and the thermodynamic catabolic energy yield on oxidation of that C (LaRowe and van Cappellin 20111; Keiluweit et al., 2016), and that energy yield is a measure of organic matter quality (Wilson and Tfaily 2018). Natural organic matter typically has NOSC values ranging from -4 to +4 with corresponding ΔG^°^_C- ox_ ranging from -54 to +174 kJ (mol C)^-1^, which suggests that most organic matter oxidation must be coupled to an energy yielding reduction in order to become thermodynamically feasible. Oxygen is capable of oxidizing compounds along the full range of NOSC values with enough energy to produce ATP. Thus, OM decomposition in the aerobic palsa is unlikely to be thermodynamically inhibited, although some evidence suggests that NOSC influences decomposability in aerobic environments as well (Graham et al., 2017). However, in the bog and fen where inundation creates anaerobic conditions and where the availability of other alternative terminal electron acceptors (such as Fe(III) or sulfate) is low, decomposition becomes thermodynamically limited, resulting in the accumulation of compounds with lower NOSC values such as fatty acids, waxes, and lignin.

While the palsa has higher oxygen availability than the other two sites, which could contribute to higher decomposition rates, the higher NOSC values of the dominant palsa plant compounds (Figure 4) are consistent with the palsa plant material also being inherently easier to decompose, regardless of the available TEAs (Keiluweit et al., 2016). The high bioavailability of palsa plants, particularly lichens (Figure 4) is contrary to generally accepted ideas that the sedges, abundant in the fen, should be the most easily biodegradable (Malmer et al 2005). The rate of litter input in the fen is highest of any of the habitats and could be faster than the microbial community can process, leading to a build-up of otherwise biologically attractive substrates (Malmer et al 2005). Similar classes of compounds accumulate in the bog and fen and both habitats have high production of condensed hydrocarbons, lignins and tannins compared to the palsa (Figure 6 B). These compounds are unlikely to be produced microbially, but are more probably due to increased (abiotic) leaching in the waterlogged bog and fen sites.

Nutrient limitation is a possible control of SOM decomposition in peatlands. While it has been shown that *Sphagnum*-dominated peatlands are nitrogen-limited (Braggazza et al., 2006), we also found evidence that dominant plants in the bog habitat are also lower in S relative to the plants from other habitats (Figure 7). This result is consistent with measurements of bulk S in the litter (Hough et al., in review) and suggests that S is limiting. In support of this hypothesis, the consumed compounds in the bog were disproportionately S-containing compounds (37% vs. 28% S in the original plant material; Figure 7). Consistent with the understanding of N limitation, the consumed compounds in the bog were also disproportionately N-containing compounds (26% vs. 17% in the original plant material; Figure 7). In contrast to the palsa, where the majority (72%) of putatively microbially produced compounds did not contain either N, S, or P (i.e., produced compounds were largely CHO-only), both the bog and fen peats had relatively large percentages of produced compounds containing these critical heteroatoms. The correlation among N and S containing compounds would be consistent with the production of microbial proteins. In other peatlands, climate effects such as warming have been associated with increases in microbial peatland cycling (Wilson et al., 2021a). Increases in the nitrogen content of decomposing peat have been observed in other studies from enhanced C losses during decomposition (Leifeld et al., 2020). The large percentages of produced compounds with N, S, and P suggest potential organic S and P cycling occurring in the anaerobic habitats.

We examined the mechanisms by which compounds are decomposed, and found that palsa has the highest overall number of transforms (i.e., potential mechanisms by which the SOM is being degraded), probably reflecting the diversity of aerobic pathways, but the fen also has a higher number of transforms compared to the bog (Figure 8). Higher numbers of transforms in the fen relative to the bog are consistent with the higher diversity of compounds in the fen plant litter stimulating microbial activity and creating a more active system. Additionally, inhibitory compounds in the bog could limit microbial activity, thereby suppressing the number of transforms utilized. In particular, the fen has a higher frequency of (de)hydrogenation (H_2_), hydroxylation (OH), demethylation followed by oxygenation (CH_2_-O), and dehydrogenation followed by oxidation (O-H_2_). Dehydrogenation and demethylation followed by oxidation (net transform: CH_4_-O) are common mechanisms of lignin decomposition (Stenson et al., 2003). Finding that these reactions are more prevalent in the fen than in the bog is consistent with the low lignin content of bog plants (i.e., *Sphagnum*) compared with the dominant fen plants (*Eriophorum*). Surprisingly, CH_4_-O is less prevalent in the palsa, where we would expect high rates of lignin decomposition due to the abundance of lignin-rich woody vegetation. Additionally, the palsa had the most diverse consumption of lignin-like compounds (Figure 6) and, in contrast to the other habitats studied, the palsa is well-oxygenated in the surface layer, which should promote the activity of the lignin-degrading enzyme phenol oxidase (Freeman et al., 2001; Sinsabaugh 2010). The FT-IR results also suggest more lignin content in the palsa plants relative to the dominant plants in the fen (Figure 1 a,c), but lower lignin content in the palsa peat relative to the fen peat, which also suggests decomposition of lignin is occurring in the fen. To reconcile these apparently conflicting results, we hypothesize that in the palsa, the greater oxygen availability allows faster, multi-step decomposition of lignin in the plant litter, such that the surface peat had already lost much of the lignin or its decomposition products; whereas in the fen, lignin decomposition is occurring (as inferred from the number of transforms), but is slowed by oxygen limitation. An alternate explanation is that although the plants are woody, litter input in any year comes mostly from the leaves, so the woody biomass has less effect on the peat.

Several transforms involving exchanges with N were important, particularly in the palsa, including oxygen or hydroxyl exchange with N, NH, or NH_3_ (Figure 8). These sorts of transforms are expected to occur when intermediates of N-fixation interact with SOM (Thorn et al., 1992, 2016; Thorn and Mikita 2000). The higher frequency of these N-involving transforms in the palsa could be related to the abundance of lichens, which are significant nitrogen-fixers in locations where herbaceous nitrogen-fixing plants are less abundant (Gunther 1989). In contrast, the mechanisms of decomposition as inferred from transform abundance in the wetter anaerobic habitats seem to be more similar to each other than either is to the drier palsa.

Mechanisms of organic matter decomposition differed between the palsa and the other habitats, but were similar between the two inundated sites suggesting that the quality of plant-derived inputs to the soil in permafrost systems influences SOM accumulation and decomposition below ground, as modified by environmental factors such as pH and oxygen availability. Shifts in plant communities in response to climate change have a profound effect on SOM composition through changing inputs. This composition in turn shapes decomposition, ultimately influencing GHG production. Nevertheless, peatlands are unique habitats in that they have a rich abundance of C but low abundance of terminal electron acceptors meaning that they are thermodynamically, yet not C, limited. Other climate forcings such as drought, which have the potential to alter the availability of TEAs, will therefore have a disproportionate influence in peatlands where an abundance of low-quality C is available for decomposition if the correct thermodynamic requirements are met.

## Acknowledgments

Funding for this research was provided by the Genomic Science Program of the United States Department of Energy Office of Biological and Environmental Research Grants DE-SC0010580 & DESC0016440. We also acknowledge funding from the National Science Foundation for the EMERGE Biology Integration Institute, NSF Award # 2022070. All data published in this manuscript is publicly accessible via the IsoGenie database https://isogenie-db.asc.ohio-state.edu/. We have no conflicts of interest to declare.

## Competing Interests Statement

The authors have no competing interests to declare.

**Supplemental Table 1:** List of Samples. Numbers indicate the number of samples analyzed by the indicated method.

|  | GPS Coordinates | FT-IR | FTICR-MS |
| --- | --- | --- | --- |
| Plants |  |  |  |
| *E. angustifolium* roots |  |  | 2 |
| *E. angustifolium* leaf |  |  | 2 |
| *E. angustifolium* (whole) |  | 1 |  |
| *A. polifolia*. roots |  |  | 2 |
| *A. polifolia*. leaf |  | 1 | 1 |
| *A. polifolia*. stem |  | 1 | 1 |
| *R. chamaemorus*. leaf |  | 1 | 2 |
| *R. chamaemorus*. stem |  | 1 | 1 |
| *R. chamaemorus*. roots |  |  | 2 |
| *E. nigrum*. roots |  | 1 | 2 |
| *E. nigrum*. leaf |  | 1 | 1 |
| *E. nigrum*. stem |  | 1 | 1 |
| *B. nana*. leaf |  |  | 1 |
| *B. nana*. stem |  | 1 | 1 |
| *B. nana*. roots |  |  | 2 |
| *S. fuscum* |  | 1 |  |
| *S. angustifolium* |  | 1 |  |
| *S. magellanicum* |  | 1 |  |
| *Sphagnum* spp |  |  | 1 |
| *E. vaginatum* roots |  |  | 2 |
| *E. vaginatum* leaf |  |  | 2 |
| *E. vaginatum* (whole) |  | 1 |  |
| Lichen |  | 1 | 2 |
| Peat |  |  |  |
| Palsa 1-5cm | N 68 21.1959 E 019 02.7974 | 1 | 3 |
| Palsa 10-14cm | N 68 21.1959 E 019 02.7974 | 1 |  |
| Bog 1-5cm | N 68 21.1973 E 019 02.8537 | 1 |  |
| Bog 10-14cm | N 68 21.1973 E 019 02.8537 | 1 | 1 |
| Fen 1-5cm | N 68 21.1992 E 019 02.8063 | 1 | 1 |
| Fen 10-14cm | N 68 21.1992 E 019 02.8063 | 1 | 1 |

**Supplemental Table 2:**
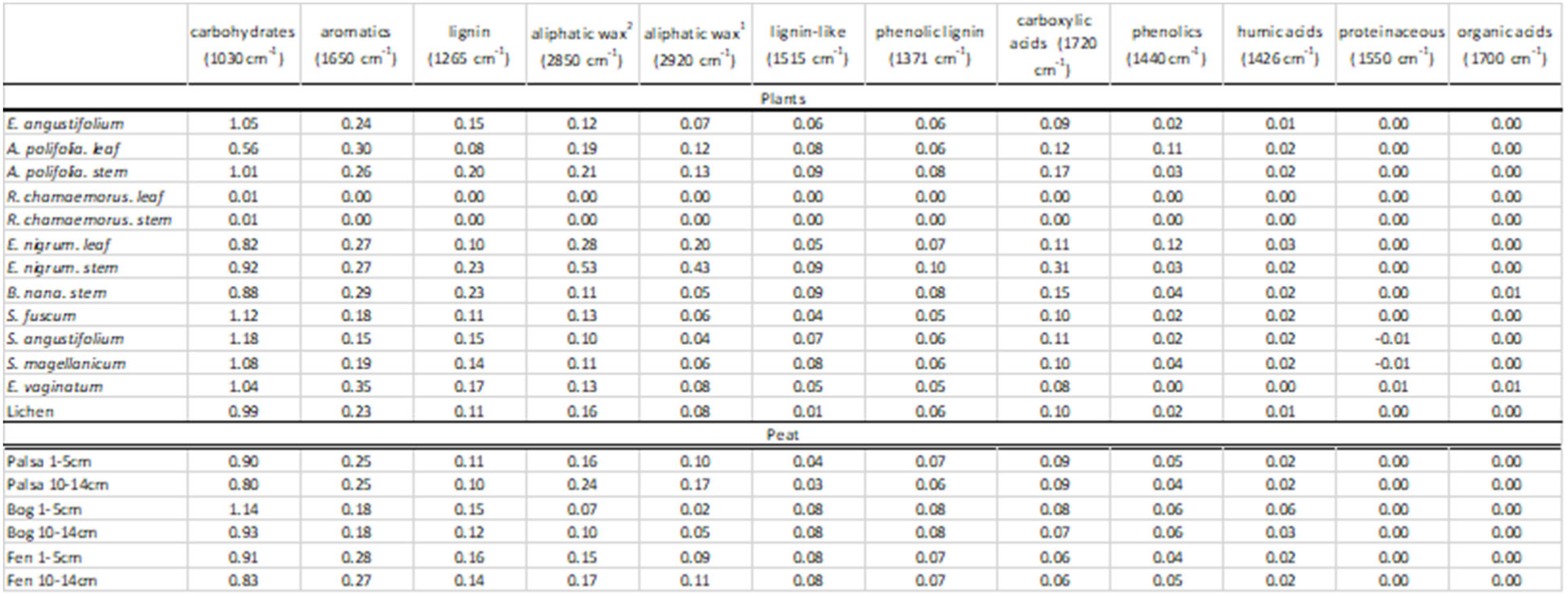
Normalized, baseline-corrected peak heights of major functionalities in the FT-IR spectra of the plants and peat from the different habitats. Peak heights are calculated according to modifications of Hodgkins et al., (2018) and reported as absorbance × 10^3^. Wavenumber assignment of functionalities is based on that given in Palozzi and Lindo (2017).

**Supplemental Figure 1:**
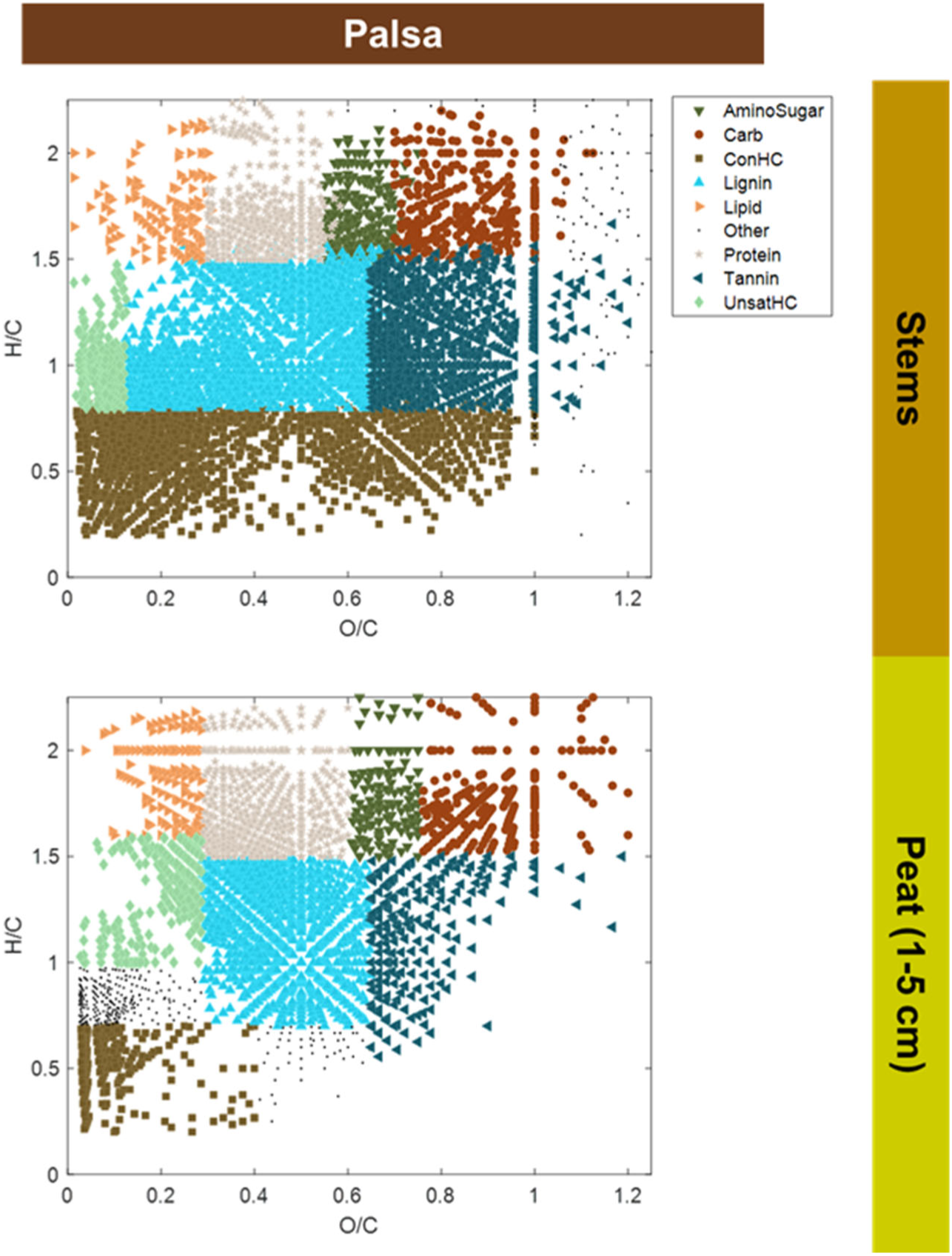
Comparing compounds observed in the plant stem extracts from palsa to those in the palsa peat. Symbols are color-coded according to major chemical classes as inferred from the FTICR-MS-derived molecular formulae. Stems were only analyzed in the palsa so bog and fen are not included.

**Supplemental Figure 2:**
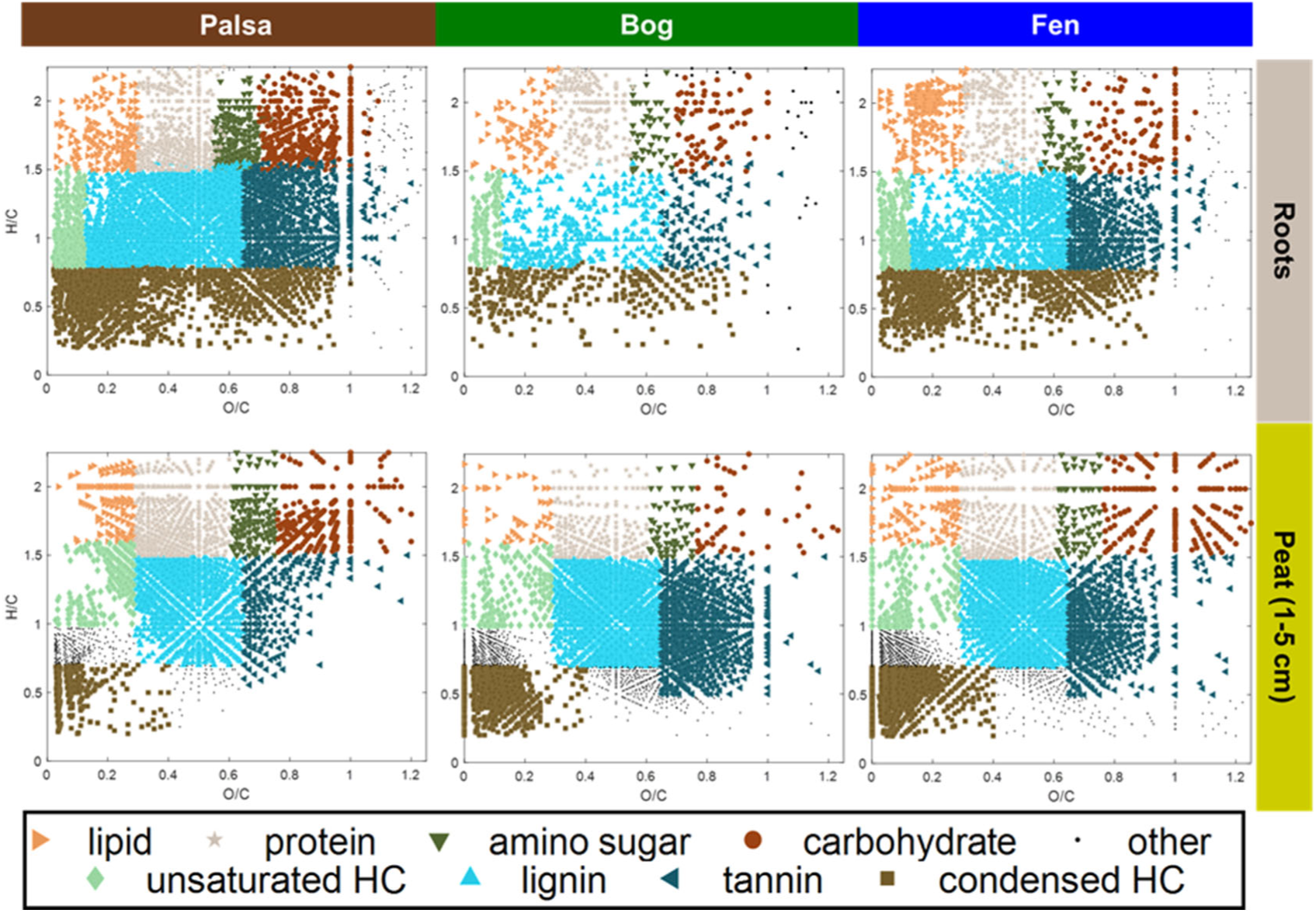
Comparing compounds observed in the root extracts from each habitat to those in the peat from each environment. Symbols are color-coded according to major chemical classes as inferred from the FTICR-MS-derived molecular formulae.

